# Paradigms of convergent evolution in enzymes

**DOI:** 10.1101/2024.04.08.588552

**Authors:** Ioannis G. Riziotis, Jenny C. Kafas, Gabriel Ong, Neera Borkakoti, Antonio J.M. Ribeiro, Janet M. Thornton

## Abstract

There are many occurrences of enzymes catalysing the same reaction but having significantly different structures. Leveraging the comprehensive information on enzymes stored in the Mechanism and Catalytic Site Atlas (M-CSA), we present a collection of 38 cases for which there is sufficient evidence of functional convergence without an evolutionary link. For each case, we compare enzymes which have identical Enzyme Commission numbers (i.e. catalyse the same reaction), but different identifiers in the CATH data resource (i.e. different folds). We focus on similarities between their sequence, structure, active site geometry, cofactors and catalytic mechanism. These features are then assessed to evaluate whether all the evidence on these structurally diverse proteins supports their independent evolution to catalyse the same chemical reaction. Our approach combines literature information with knowledge-based computational resources from, amongst others, M-CSA, PDBe and PDBsum, supported by tailor made software to explore active site structure and assess mechanism similarity. We find that there are multiple varieties of convergent functional evolution observed to date and it is necessary to investigate sequence, structure, active site geometry and enzyme mechanisms to describe such convergence accurately.

## Introduction

Nature shapes biological macromolecules during evolution, allowing mutable elements to change, or conserving them by natural selection. The extent of selective pressure is variable, and this contributes to functional divergence in proteins of common ancestry. We have recently reviewed some of the concepts in enzyme evolution, especially functional divergence from a mechanistic viewpoint[1]. All known enzyme reactions are performed by a relatively limited number of structural folds. However, the chemical reaction space is vast and there may be many biological reactions we have yet to discover. Despite this, amongst the many reactions we know, many are catalysed by more than one family of enzymes, that are not related by evolution. Surprisingly, this is quite common, with a reaction on average being catalysed by ∼2 evolutionarily unrelated enzymes[2,3]. Such analogues are called isofunctional enzymes or isozymes[4] and the evolutionary phenomenon that describes them is convergent evolution[5].

It is not uncommon for a catalytic process, or part of it, to be facilitated by a set of chemical groups in a well-defined geometry that occur in multiple enzymes[6]. These 3D constellations often drive a unique sequence of events of the catalytic mechanism. Popular examples are the various catalytic triads in proteinases[7,8] that hydrolyse peptide bonds by nucleophilic substitution. Although the nucleophile position might be occupied by different residues, usually Ser or Cys, the spatial arrangement of the triad (Nucleophile-His-Acid) is highly similar among most proteinases and has evolved independently in analogues[9]. In all cases, the same sequence of catalytic steps takes place. Similarly, the binding affinity for a substrate depends on the geometry of residues in the binding pocket, where any conformational changes can dramatically affect ligand selectivity. In the case of small ligands (especially metal ions), binding constraints can be strong, leading to a common geometry in unrelated proteins (e.g. Fe-SO metal cluster binding site).

In the context of convergent evolution, we can ask questions such as:

1. Since functional convergence can occur in multiple ways, what are the different facets of such convergence?
2. Can we distinguish different paradigms, and which occur most often in nature?
3. Why has nature evolved the same function more than once?

Leveraging the integrated information within the Mechanism and Catalytic Site Atlas (M-CSA) –i.e. sequence, structure, homologues, functional annotation of residues, curated annotation of catalytic residues and explicit description of the mechanism – herein we aim to address these questions and define broad paradigms of enzyme convergent evolution.

We present 38 detailed examples of isozymes, characterising their catalytic sites and mechanisms to illustrate convergent evolution of function. We also provide evidence that categorisation into fine-grained paradigms is not straightforward, and in several cases, enzymes adhere to multiple paradigms. Our observations highlight some limitations in enzyme function and structure classification systems, and the need to consider additional parameters such as local active site geometry, cofactor selectivity and chemistry to classify ambiguous evolutionary relationships. This systematic analysis of multiple enzyme features (sequence, local and global structure, catalysed reaction, catalytic mechanism, substrate selectivity, promiscuity etc.), provides extra knowledge, useful for enzyme design by computational methods and directed evolution[10].

## Results

The Mechanism and Catalytic Site Atlas (M-CSA) aims to include entries for all known enzyme reactions for which sequence, structure and functional data are available, and so inevitably the content reflects the protein coverage in UniProt and PDB, which are both biased towards well studied enzymes. The rationale of M-CSA content and curation is broadly based on including one reference entry for every unique EC number, supplemented by enzymes with the same EC number but with a different mechanism. We took advantage of this redundancy to identify and compare pairs of enzymes, which appear to perform the same reaction but belong to different structural families (i.e. examples of convergent evolution). To this end, we extract two datasets of paired enzymes: 1) all pairs of entries in M-CSA (2021 update) with the same EC number at reaction level (sub-sub class - EC x.x.x.-), and 2) all pairs of entries with the same EC number performing the same reaction on the same substrate (sub-sub-subclass - EC x.x.x.x). In a pair, the two enzymes might be homologues (their catalytic domains belonging to the same CATH superfamily) or analogues (when the proteins are from different CATH superfamilies, thus are likely to have evolved independently). The whole dataset consisted of 6,345 pairs, many of which occuring more than once, so we further filtered for unique EC pairs, reducing this number to 2,209. Using protein CATH numbers as filters we distinguished homologous pairs (identical CATH number) from analogous pairs (different CATH numbers). These results are presented in Figure 1, which compares the two datasets.

**Figure 1:**
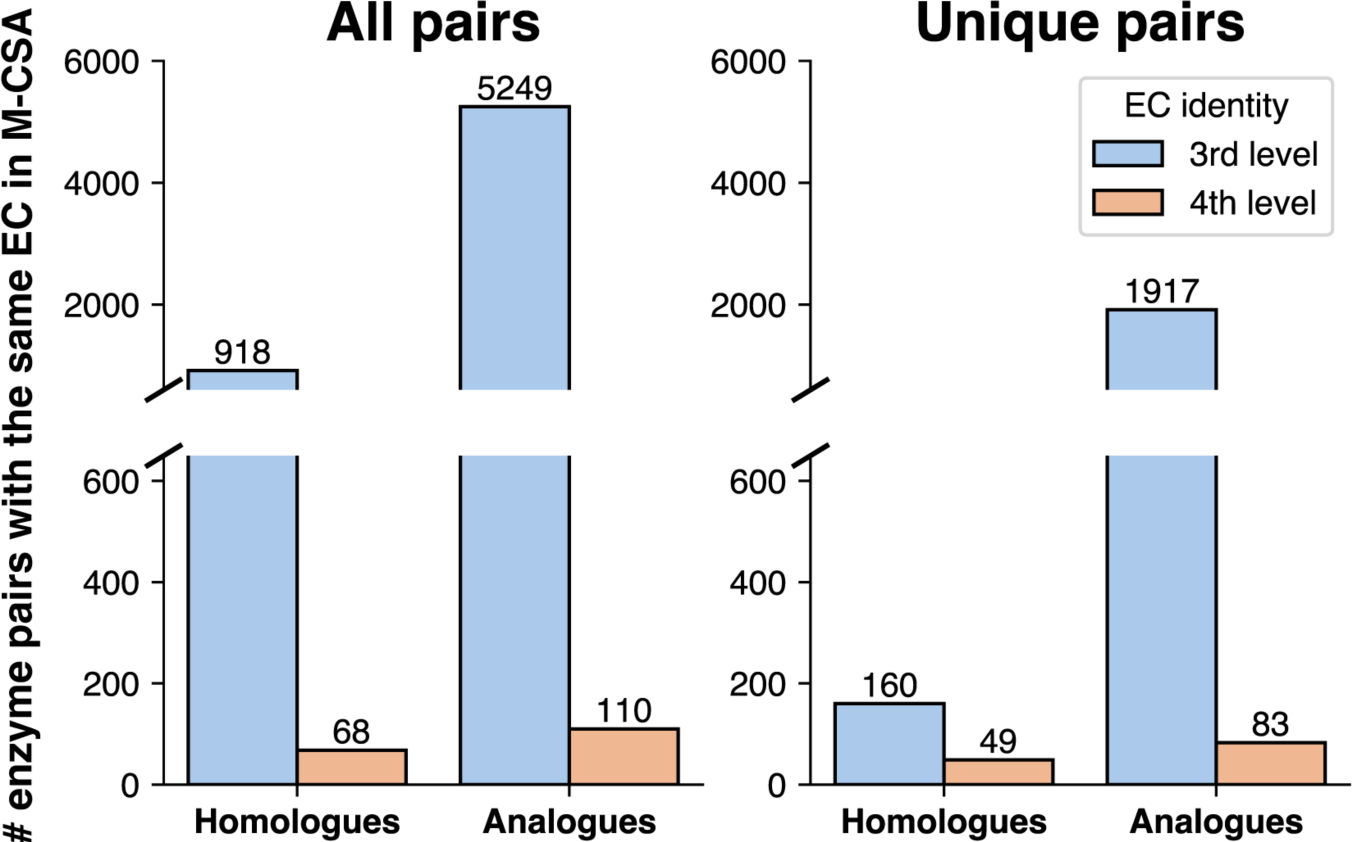
Number of enzyme pairs in M-CSA sharing the same EC number, when looking at the sub-subclass (3^rd^ number) or sub-sub-subclass (4^th^ number) level. Homologues have identical catalytic domain CATH numbers and analogues have different CATH numbers.

We find many more analogous pairs than homologues (only 9-15% are homologues, covering 41 EC numbers in total) reflecting the content of the M-CSA. Most of the homologous functional duplicates differ in their mechanism, and a few are attributed to curation mistakes. We also find that considering only the reaction at the third level (i.e. ignoring substrate specificity by removing the 4^th^ EC level filter) leads to a steep increase of the number of enzyme pairs. This increase remains significant even when redundancy is removed (All vs Unique pairs in Figure 1). However, we already know that in nature, most new functions evolve from old functions[11] by simply changing the substrate – but here we are deliberately targeting pairs of enzymes with the same function but in different structural families – i.e. analogues) Out of these sets, we selected the unique analogous pairs matching at the 4^th^ EC level (83 pairs in total, with both members of the pair catalysing the same reaction with identical substrates). After filtering (see Methods) we ended up with 38 pairs, to analyse and exemplify three paradigms of convergence (Figure 2).

**Figure 2:**
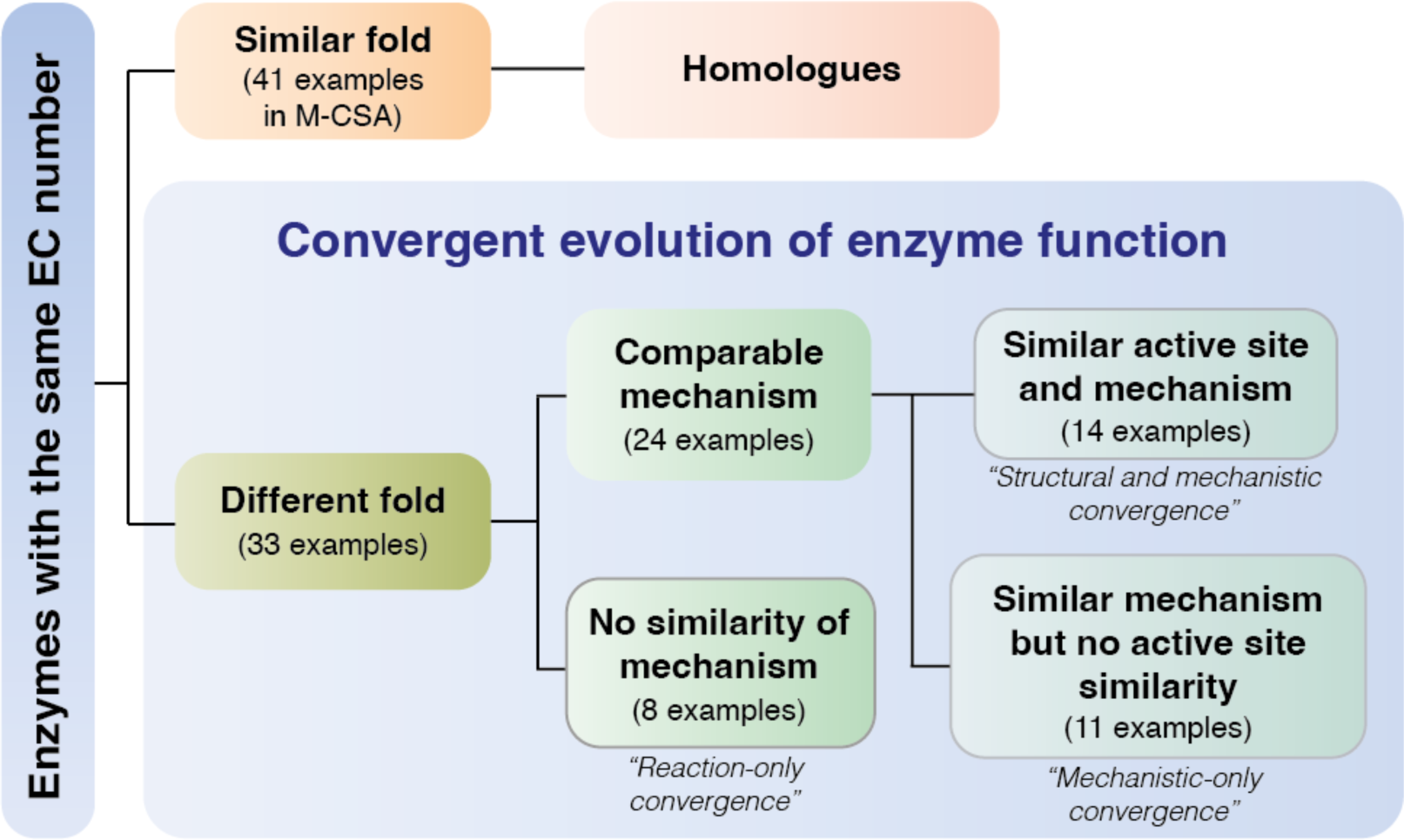
Evolutionary relationships of enzymes of the same function. Paradigms of convergence are quoted.

These 38 pairs were examined for similarities in different facets, some of which we discussed in the context of bioinformatics in our recent review[1]. Using computational tools and/or literature reviewing, we looked at the following:

1. overall structural similarity (CATH superfamily assignment and 3D fitting)
2. active site structural similarity (quantified by RMSD of aligned residues)
3. mechanistic features (catalytic steps[12,13] or/and residue roles)
4. substrate similarity
5. cofactor presence/similarity

Our results are organised as pairs or triplets of analogues in Table 1-3. Three selection rules are implicit in this survey of convergent evolution: 1) catalytic domains should always belong to different CATH superfamilies, 2) All 4 E.C. numbers should be identical and 3) isozymes should share at least one identical product and substrate. This means that different paradigms of evolutionary convergence are defined by similarities in the active site composition and geometry, catalytic mechanism and cofactor selectivity. Focussing solely on mechanism, three general ‘cases’ are observed. Firstly, analogues that are completely different in their mechanism of catalysis. In the second ‘case’, analogues that are similar but only part of the reaction is the same (e.g. phospholipases (EC 3.1.1.4) had similarities in the steps 1 & 2 of one enzyme and steps 3 & 4 of the other respectively). In the 3^rd^ case, mechanisms between the analogues are essentially the same, with only minor differences. For these, the mechanisms are nearly identical except for differences in the catalytic residues being used (e.g. both analogous Papain-like cysteine proteases (EC 3.4.22.28) use a Cys-containing catalytic triad, but the His activator can be Asp or Asn). Combining these mechanistic paradigms with any localised structural similarities in the active site[14], led us to define three general paradigms of convergence in isozymes, described in the following section.

**Table 1:** Isoenzyme examples with structural and mechanistic convergence.

| Enzyme | EC | CATH domain | M-CSA ID | PDB ID | Residue composition | 3D-aligned residues | RMSD (Å) | Mech. sim. (calc. perc.) | Cofactor | Notes | Refs. |
| --- | --- | --- | --- | --- | --- | --- | --- | --- | --- | --- | --- |
| Carbonate dehydratase | 4.2.1.1 (a) | 2.160.10.10 | 516 | 1QRM | EEHHHNQ | His117 Glu62 His122 His81 | 1.45 | Yes (30%) | Co<sup>3+</sup> | Similar metal binding site. Sulfate moieties bind close to HCO3 positions in both analogues. | [15,16] |
|  |  | 3.10.200.10 | 216 | 5GMN | EHHHHT | His94 Glu106 His119 His96 |  |  | Zn<sup>2+</sup> |  |  |
| Protein tyrosine phosphatase | 3.1.3.48 | 3.40.50.2300 | 462 | 2P4U | CCDNRS | Ser20 Cys13 Arg19 | 1.30 | Yes (n/a) | - | Both analogues are mammalian. Significant catalytic core 3D similarity in completely different fold contexts. Divergence followed by convergence is plausible. | [17,18] |
|  |  | 3.90.190.10 | 469 | 2GJT | CDQRT | Thr1143 Cys1136 Arg1142 |  |  | - |  |  |
| 3-hydroxydecanoyl dehydratase | 4.2.1.59 | 3.10.129.110 | 972 | 2VZ9 | DGHHLV | Any885 Asp1033 Any888 His878 | 0.62 | Yes (27%) | - | One analogue is mammalian, the other is bacterial. Significant catalytic core 3D similarity. Either a case of extreme convergence or distant homology. | [19] |
|  |  | 3.10.129.10 | 10 | 1MKA | CDGHV | Any76 Asp84 Any79 His70 |  |  | - |  |  |
| Endo-1,4-β-xylanase | 3.2.1.8 | 2.60.120.180 | 432 | 2B42 | EENYY | Glu172 Glu78 Asn35 | 1.67 | Yes (n/a) | - | Sugar is cleaved with very similar mechanisms using a Glu-Glu dyad. | [20,21] |
|  |  | 3.20.20.80 | 548 | 1FHD | DEEHN | Glu233 Glu127 Asn169 |  |  | - |  |  |
| Peptidylprolyl isomerase | 5.2.1.8 | 2.40.100.10 | 189 | 1XO7 | FFHLNQR | Any103 Phe62 Phe114 | 2.56 | Yes (n/a) | - | Two Phe residues form a hydrophobic pocket for substrate side chain rotation. Main chain residue is H-bonded to substrate carbonyl. Both human-derived, but 362 induces reversible protein folding. | [22,23] |
|  |  | 3.10.50.40 | 362 | 1NSG | DDFIYY | Any56 Phe36 Phe99 |  |  | - |  |  |
| Lysozyme | 3.2.1.17 | 1.10.530.10 | 203 | 3IJU | DDENNS | Glu35 Asp52 Asp48 | 2.65 | Yes (27%) | - | Hydrolysis mechanism is similar, with slightly different spatial arrangement of residues. | [24,25] |
|  |  | 3.20.20.80 | 774 | 1H09 | DDDE | Glu94 Asp92 Asp10 |  |  | - |  |  |
| Protein-methionine-S-oxide reductase | 1.8.4.11 | 3.30.1060.10 | 122 | 4D7L | CCCDEYY | Asp130 Cys52 Cys206 | 3.14 | Yes (23%) | - | Cys residues have slightly different roles in the two enzymes, but two catalytic steps are similar. | [26,27] |
|  |  | 2.170.150.20 | 715 | 1L1D | CCDHHR | Asp484 Cys440 Cys495 |  |  | - |  |  |
| Ubiquitinyl hydrolase 1 | 3.4.19.12 | 3.40.532.10 | 597 | 4I6N | CDHQ | Any243 His340 | 1.18 | Yes (n/a) | - | Common Cys deprotonation by His with Cys also contributing its backbone in oxyanion hole | [28,29] |
|  |  | 2.40.10.10 | 830 | 2GZ9 | CEGH | Any147 His40 |  |  | - |  |  |
| Papain-like cysteine protease | 3.4.22.28 | 3.90.70.10 | 805 | 1Y4H | CHNQ | Any243 His340 | 1.26 | Yes (n/a) | - | Very similar active sites, oxyanion hole is Cys backbone and Ala in one analogue and Cys backbone and Asn in the other. Asn and Glu modify the pKa of the His in both. | [30,31] |
|  |  | 2.40.10.10 | 477 | 6FFS | CEGH | Any147 His40 |  |  | - |  |  |
| Ornithine decarboxylase | 4.1.1.17 | 3.40.640.10 | 860 | 1ORD | DHK | Lys355 His223 | 1.75 | Yes (n/a) | PLP | Identical mechanism between bacterial/mammalian analogues. Their only difference is the mirrored arrangement of residues. | [32,33] |
|  |  | 3.20.20.10 | 937 | 5GJO | EHK | Lys51 His179 |  |  | PLP |  |  |
| Phosphohistidine-D-mannose phosphotransferase | 2.7.1.191 | 1.20.58.80 | 514 | 1E2A | HHQ | His78 His82 | 0.44 | Yes (n/a) | - | Two His are at almost identical positions, but reside on different secondary structure elements. Roles in mechanism are similar. | [34,35] |
|  |  | 2.70.70.10 | 513 | 3OUR | HHT | His91 His76 |  |  | - |  |  |
| Cytosine deaminase | 3.5.4.1 | 3.20.20.140 | 710 | 1K6W | DEHHHQ | His214 Glu217 | 0.69 | Yes (35%) | Zn<sup>2+</sup> | In both analogues, His ligates a metal. Glu deprotonates a water molecule and protonates cytosine N3. Difference lies in the combination of metal-ligating residues. | [36,37] |
|  |  | 3.40.140.10 | 636 | 1UAQ | CCEHS | His62 Glu64 |  |  | Zn<sup>2+</sup> |  |  |
| Endonuclease | 3.1.21.1 | 3.90.540.10 | 791 | 2GZI | EHHHHRR | His103 His127 Glu100 | 1.43 | Yes (30%) | Zn<sup>2+</sup> | Similar initial steps of nucleophilic attack on phosphate via a proton relay. A Glu-His dyad is present in both analogues. | [38] |
|  |  | 3.60.10.10 | 41 | 2A40 | DDEEHY | His252 His134 Glu78 |  |  | Mg<sup>2+</sup> |  |  |
| 3-dehydroquinate dehydratase | 4.2.1.10 | 3.40.50.9100 | 55 | 4KI7 | EGHNNPRRY | Glu99 His101 | 0.83 | No (0%) | - | Glu-His proton relay motif present in both analogues. Glu activates His, but the overall mechanistic context is different. | [39,40] |
|  |  | 3.20.20.70 | 54 | 1QFE | EHK | Glu86 His143 |  |  | - |  |  |

### Paradigms of convergent evolution

38 cases of isozymes in M-CSA reveal three paradigms of functional convergence:

1. **Structural and mechanistic (Table 1).** Two or more catalytic residues align in 3D, and there are similarities in the mechanism. This is a very common paradigm, with 14 examples. In some cases, a subset of catalytic residues may align in 3D, but the mechanisms are nevertheless different. This sub-paradigm is not common here (2 examples), and active site similarities, if found, are usually located on a ubiquitous motif (e.g. an activator stabilised by an acid or a common motif binding a catalytic metal).
2. **Mechanistic-only (Table 2).** Similarities in mechanism are observed, with no aligned catalytic residues. This is also a very common paradigm (11 examples). In such cases, a general similarity in the active site might exist, but this is not detectable in superposition.
3. **Reaction-only (Table 3).** Only the overall reaction is similar, with no similarities in mechanism or active site geometry. A moderately common paradigm for which 8 examples are found.

**Table 2:**
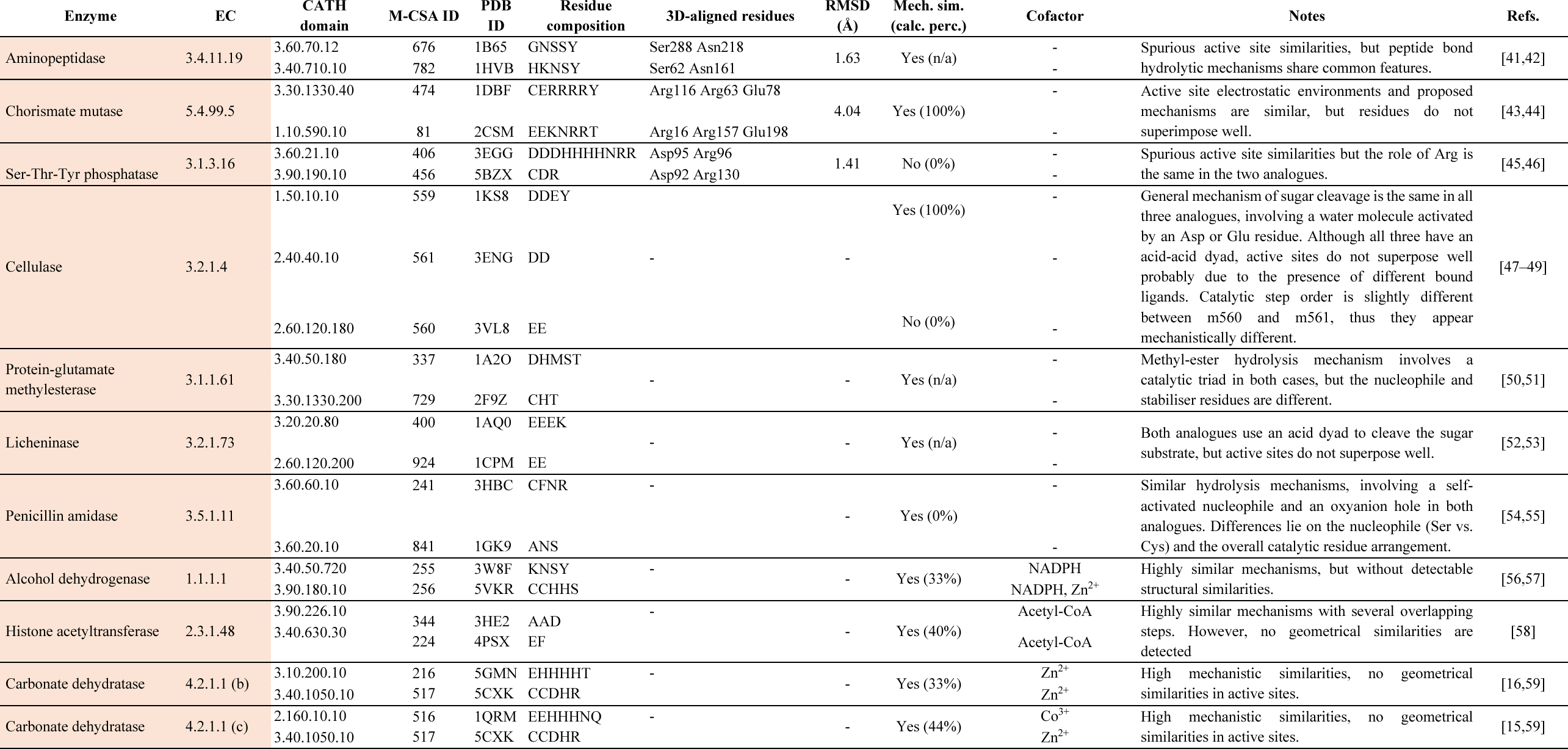
Isoenzyme examples with mechanistic-only convergence.

**Table 3:**
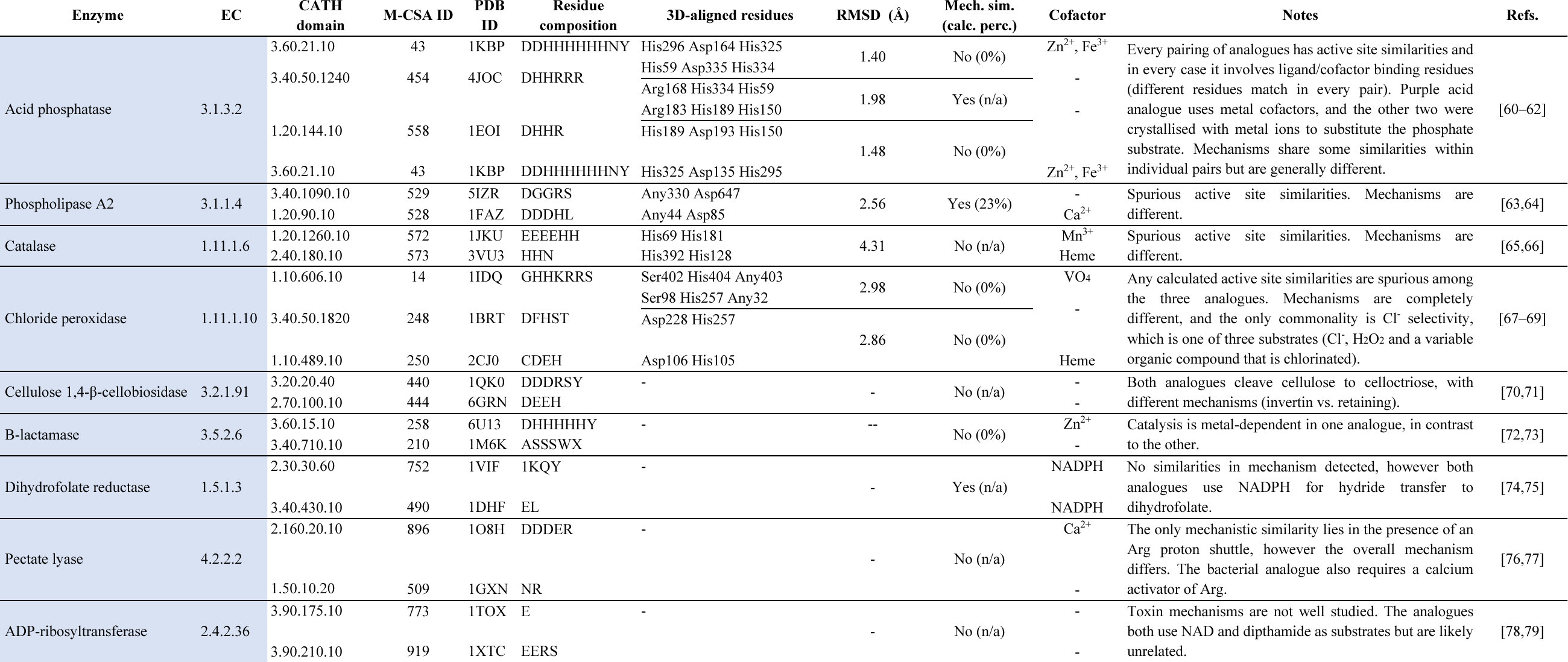
Isoenzyme examples with reaction-only convergence.

Paradigms can be divided further according to cofactor selectivity: a) Similar cofactors are used and b) different cofactors are used, or one analogue uses a cofactor while the other does not.

There are also cases in this dataset that could be assigned to more than one paradigm or sub-paradigm. For instance, the two unrelated 3-dehydroquinate dehydratase analogues (EC 4.2.1.10) use a His-Glu dyad, where Glu as an activator of His, but these occur in different mechanistic contexts. Chorismate mutases (EC 5.4.99.5) are also ambiguous and can be assigned to either paradigm 1 or 2, depending on how structural and functional similarities are interpreted.

### Active site 3D similarities accompany mechanistic similarities (Table 1)

Several examples in our data correspond to mechanistic analogues, with similarities in mechanism being reflected in three dimensions (structural and mechanistic convergence).

To illustrate this paradigm, we selected the human and yeast carbonate dehydratase analogues. These classes of carbonate dehydratases (alpha and gamma) have converged to catalyse the reversible conversion of carbon dioxide and water into hydrogencarbonate (Figure 3). They use a similar mechanism where the most important catalytic step is a nucleophilic attack on the carbon dioxide by a hydroxide ion, which is being stabilized by a positively charged metal ion (mechanism step picture in Figure 3). Interestingly, although the enzymes use a different metal ion, Co^3+^ in the case of the gamma class and Zn^2+^ for the alpha class, both active sites provide three histidine residues to hold the metal ion in place. While the coordination sphere of Zn^2+^ is satisfied by the three histidines and the hydroxide, Co^3+^ binds an additional water molecule and as the reaction proceeds, the hydrogencarbonate product. The active sites also differ in the residue that deprotonates the water molecule to generate the hydroxide (a His for alpha vs. a Glu for gamma) and the residues that orient the carbon dioxide (Glu and Thr for alpha class vs. Asn for gamma class).

**Figure 3:**
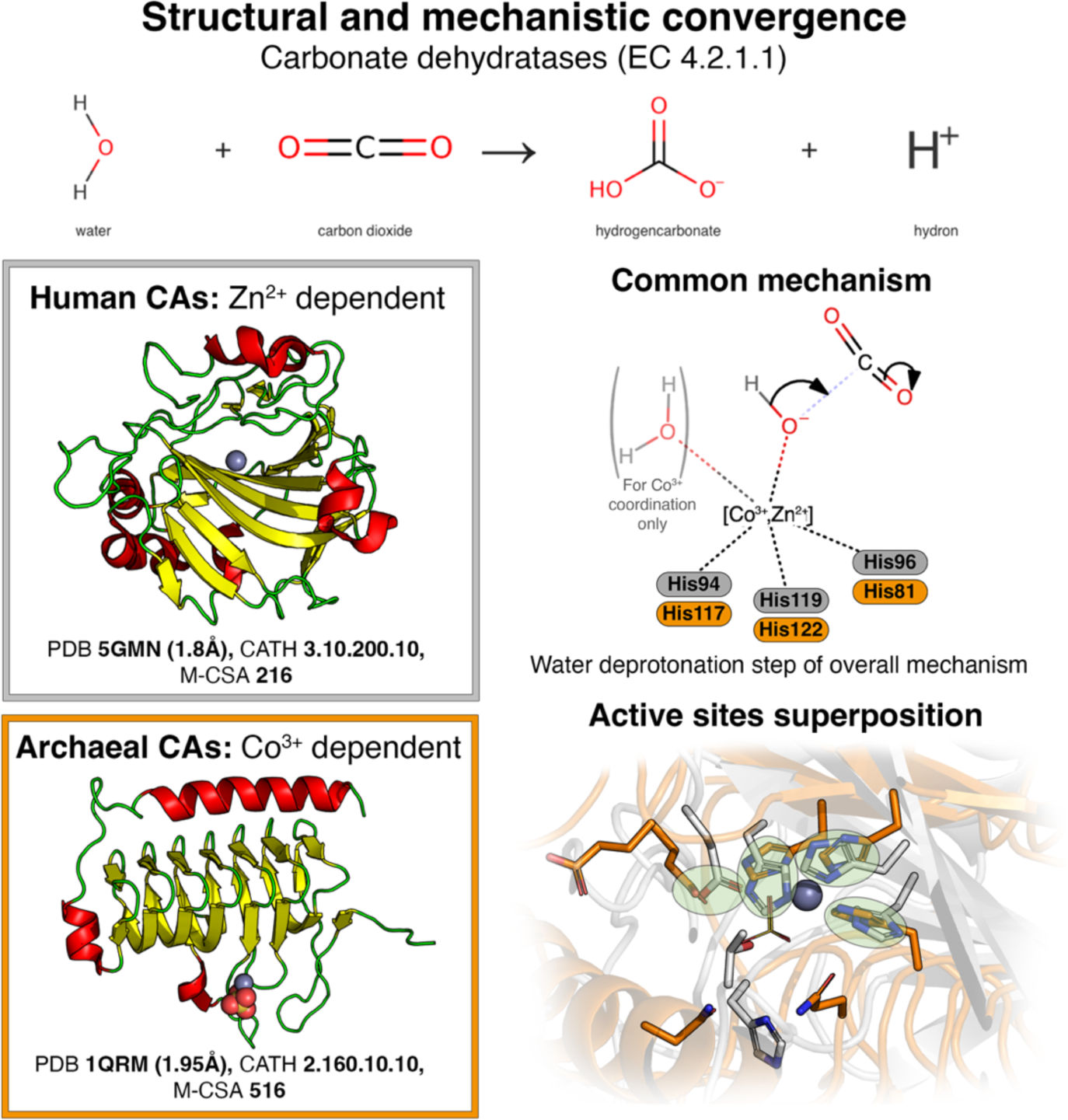
Structural and mechanistic convergence between human (grey) and archaeal (orange) carbonate dehydratases. Green circles in the active sites superposition indicate pairs of 3D-aligned catalytic residues.

Another illustrative example is the pair of mammalian/bacterial ornithine decarboxylases (EC 4.1.1.7) that both use an (Asp/Glu)-His-Lys residue triad to cleave a carboxyl group from the ornithinium substrate (Figure 4a). A PLP cofactor is involved in both mechanisms, that is initially covalently bonded to the catalytic Lys. Global sequence alignment showed low similarity although the local alignment has relatively high coverage. The enzymes have no overall structural similarity, belonging only to the same primary CATH class (1^st^ level). In their active sites, the two analogues have mirrored but similar arrangement of catalytic residues, with Lys and His endpoint atoms superposing at RMSD 1.75Å. By these observations, homology is unlikely; instead, convergence in function, local structure and mechanism is inferred.

**Figure 4:**
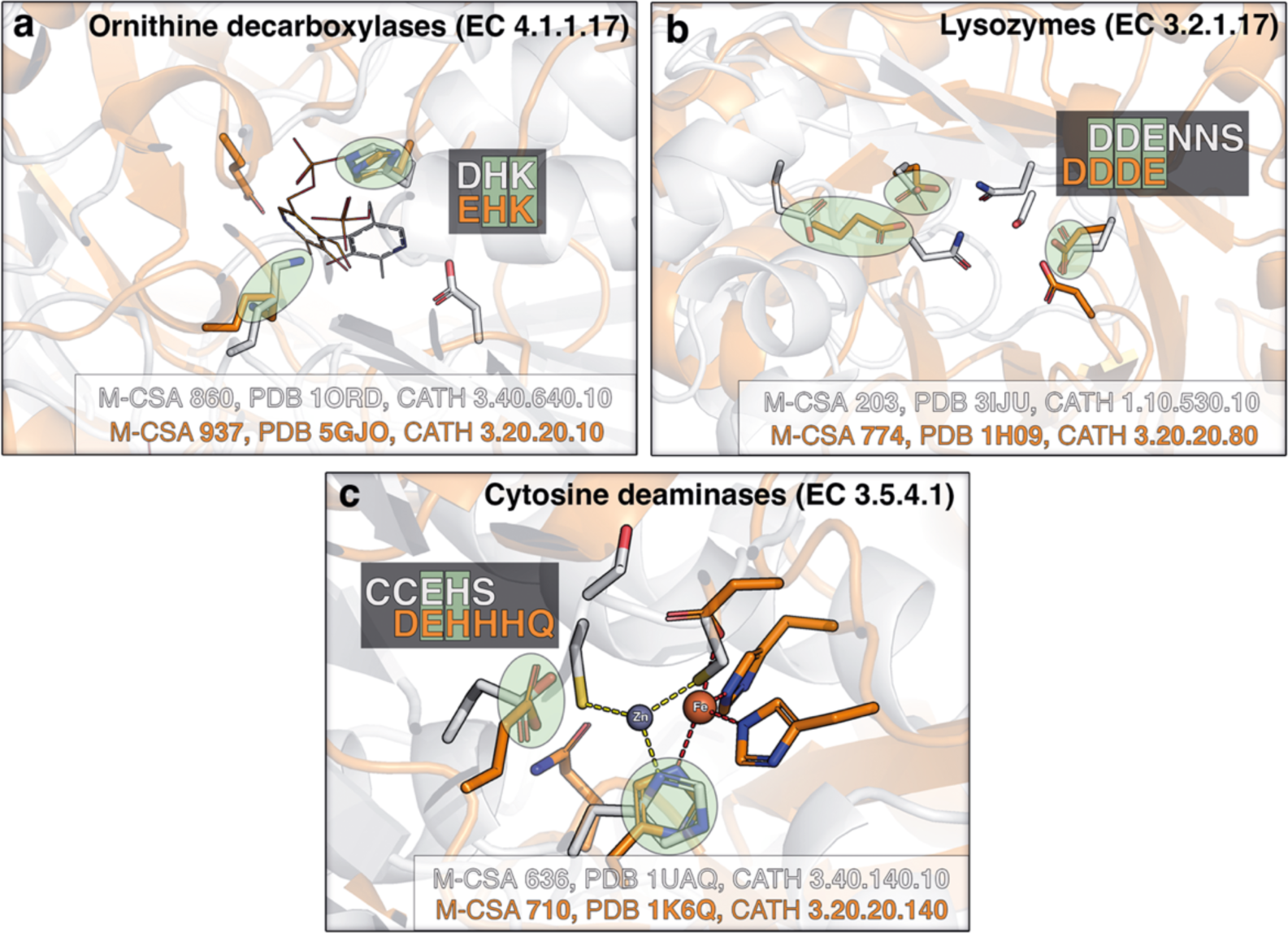
Active site similarities in three pairs of analogues with structural and mechanistic convergence. a) Bacterial (grey) and mammalian (orange) ornithine decarboxylases. b) Chicken (grey) and viral (orange) lysozymes. c) Yeast (grey) and bacterial (orange) cytosine deaminases. Green circles indicate 3D-aligned catalytic residue pairs. Residue alignment is also shown on the active site residue composition (pseudo-sequence as in Table 1) with the aligned residues highlighted in green.

Structural constraints may be relaxed in some cases, with slightly different functional atom 3D configurations facilitating the same mechanistic sequence. Mammalian and viral lysozyme analogues (EC 3.2.1.17) represent this paradigm variant (Figure 4b), between which the carbonyl groups of three acidic residues (one Glu and two Asp) loosely superpose at RMSD 2.65Å. Glycosyl hydrolases GH22 (viral) and GH25 (mammalian) break down β-MurNAc-(1- >4)-β-D-GlcNAc to N-acetyl-β-D-muramic acid and N-acetyl-β-D-glucosamine via hydrolysis, with two proposed mechanisms in each analogue. They have virtually no structural similarity, and GH25 uses a choline ion to facilitate cell wall binding, which is not directly involved in catalysis but does increase catalytic activity, while GH22 does not use any cofactors. Global and local sequence alignments showed low similarity, and except for Glu35 in GH22 lining up with a non-catalytic Glu in GH25 in the global alignment, no active site residues line up with an identical (or near-identical) residue in any alignment. The only differences between them are in which atoms serve as donors/acceptors, and in one case the active site regeneration chemistry is different. Every proposed mechanism directly involves a Glu with an intramolecular O atom (the latter is present in three out of total four proposals) to break the glycosidic bond. Both have the presence of two acidic residues hugging the substrate, similar to cellulases. The analogues show 27% mechanistic similarity, and given the low sequence and structural similarity, it is likely that they evolved this function separately.

Evolution can also shape active sites in a modular way, with resulting enzymes having similarities in the mechanism, but differing at a technicality. A good example is cytosine deaminases (EC 3.5.4.1) where both analogues use a catalytic metal, coordinated by different residues sets (Figure 4c): 3 His and an Asp binding a Fe ion in the yeast enzyme, while 1 His and 2 Cys bind a Zn ion in its bacterial analogue. In both cases, the metal binding His and the proton donor/acceptor Glu have similar 3D arrangement, with the latter protonating the cytosine N3 by obtaining a proton from a water molecule.

Within this paradigm, we also distinguish analogue pairs that can be characterised alternatively, by a special case we name as “structural-only convergence”. This can be defined as structural convergence in the active site, but with similarities only observable in small motifs such as catalytic metal binding sites or activator-stabiliser dyads (e.g. His-Glu or His-Asp). This is clearly seen in two examples that both contain a His-Acid dyad: 3-dehydroquinate dehydratases (EC 4.2.1.10) and endonucleases (EC 3.1.21.1). In all four enzymes, the acid (Asp/Glu) interacts with a His via an H-bond, to activate and/or sterically orientate it, however in different mechanistic contexts. The same dyad is also present in catalytic triads of Ser/Cys proteases with the acid playing the same role.

### Mechanistic similarities with different residue toolkits (Table 2)

We observed cases where mechanistic similarities might exist, without detectable active site similarities (mechanistic-only convergence). In such cases, any structurally aligned residues found by our alignment algorithm are most likely spurious or superpose loosely. This is demonstrated in fungal and bacterial chorismate mutases (EC 5.4.99.5), between which 3 catalytic residues superpose at a very high RMSD (4.04Å – considered dissimilar), but the overall electrostatic environment that facilitates isomerization of the chorismate substrate is similar, after manual inspection.

A representative example for the “mechanistic-only” paradigm is licheninases (EC 3.2.1.73) produced by both plant and bacteria for cell wall degradation. These are unrelated in primary sequence (20% sequence identity) and tertiary structure (Figure 5). Despite the differences in protein fold and local geometry they retain a similar mechanism for cleaving cereal β-D glucans and lichenin with the formation of an intermediate covalent bond between the substrate and a glutamic acid in the enzyme during catalysis[80].

**Figure 5:**
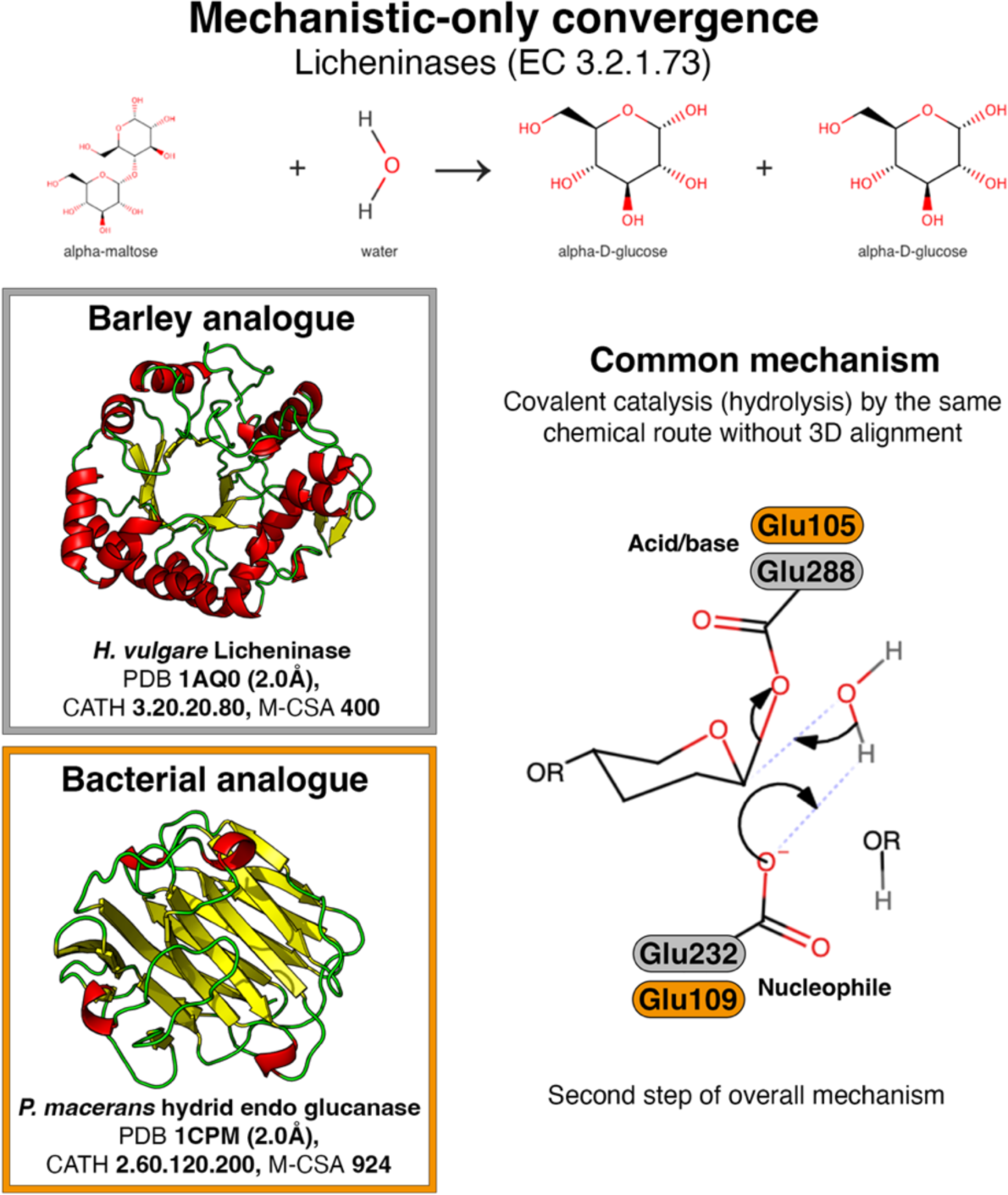
Mechanistic-only convergence between plant (grey) and bacterial (orange) licheninases.

Aminopeptidase isoenzymes from *Ochrobacterium anthropi* (EC 3.4.11.19) also conform to this paradigm. They are also a great example of a duplicate function within the same organism/cell/compartment, presenting the evolutionary question of why two isoenzymes would co-exist. The two analogues are known as DmpA (M-CSA entry 676) and DmpB (M-CSA entry 782) with no evidence of a common ancestor between them. However, their mechanisms are similar without geometrical convergence in the active site. DmpA is poorly characterised, mostly because it is expressed in low quantities[81]. DmpB is more abundant and better characterised, and is also an ancestor of penicillin resistance conferring enzymes (β-lactamase activity)[42]. It is plausible that the two analogues co-exist in the same organism, because they are regulated differently. DmpA is also functionally promiscuous, with D-esterasic and D-amidasic activities as well as autoproteolysis[43], suggesting that aminopeptidase activity is not the primary one, compared to DmpB, whose function is more specific.

### Convergent evolution with different residue toolkits and mechanism (Table 3)

Enzymes can converge in the overall catalysed reaction, with different stepwise chemistries, with no or limited similarities (reaction-only convergence – no mechanism or structure similarity). Reaction-only convergence is moderately abundant in M-CSA (9 examples) and refers to enzymes that have the same EC number (i.e. they catalyse the same reaction) but have no detectable similarities in their 3D structure. These cases (Table 3) are particularly interesting when enzyme analogues of different CATH families catalyse the same reaction in a given phylum.

In the case of fungi for example, the two cellobiosidases (EC 3.2.1.91) act on the cellotetraose substrate using a different ensemble of residues and mechanisms (inverting vs. retaining[82]) for bond cleavage, giving isomeric products of opposite stereochemistry[83]. Similarly, structurally different β-lactamases (EC 3.5.2.6) in bacteria perform the same catalytic function using either a protein-water or a distinct protein-metal mediated process (Figure 6). Reaction-only convergence can also be observed across taxa, for example the peroxidases (EC 1.11.1.10) from fungus and bacteria (Table 3). In this case, the oxidation of the substrate is achieved by utilising significantly different mechanisms through the selective exploitation of non-proteinous components like Chlorine, Heme or Vanadium.

**Figure 6:**
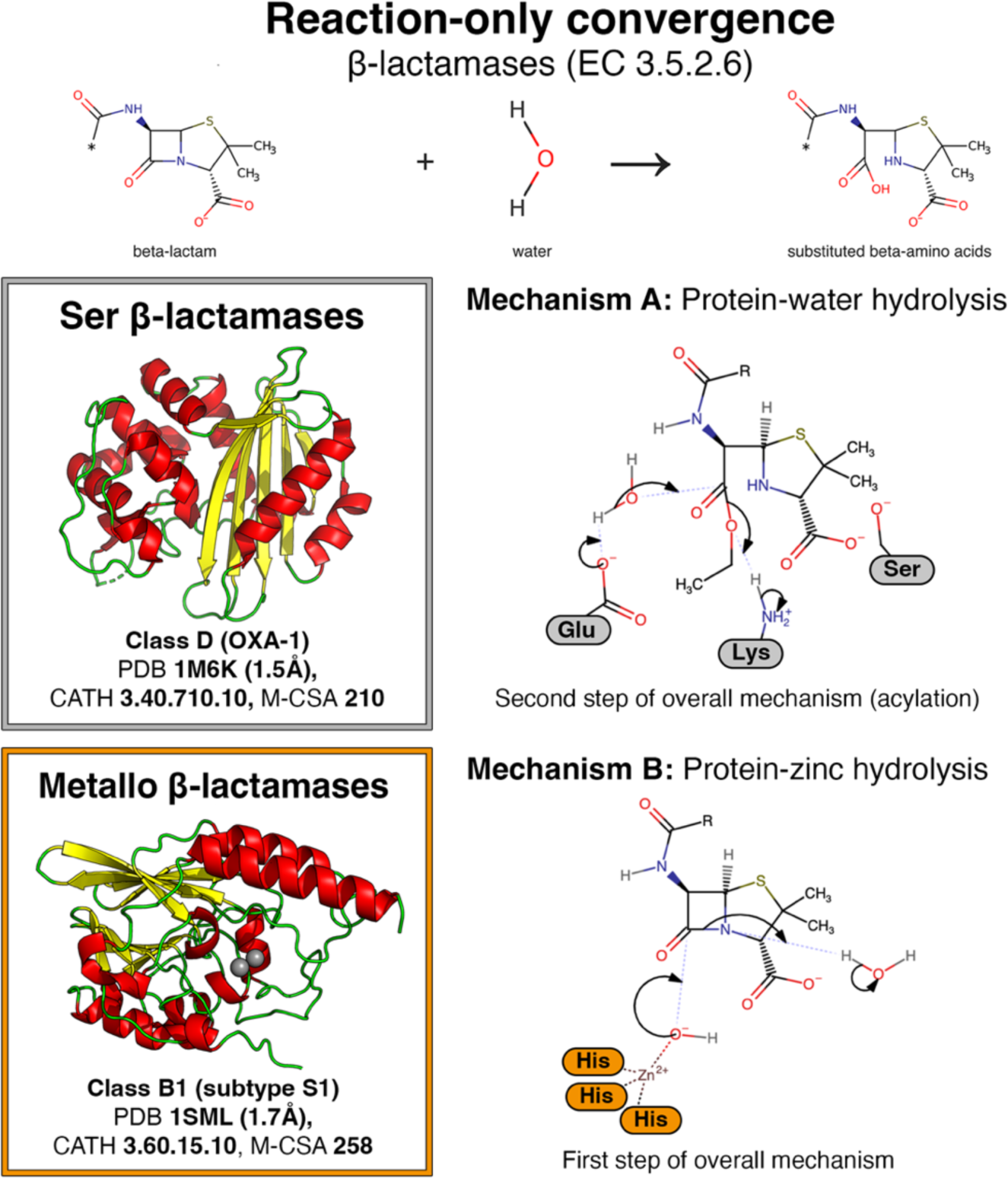
Reaction-only convergence in β-lactamases. The exact acid and bases, and the sequence of catalytic events vary in different Ser β-lactamases. Metallo β-lactamases bind two metal ions in the active site, but only the one participating in the described steps is shown in the mechanism diagram.

### Convergence and distant divergence are often indistinguishable

Convergence is not easily distinguished from distant divergence, especially in the case of generic metabolic enzymatic activities. Such isoenzymes have little sequence and overall structural similarity, belonging to different CATH superfamilies (H level) or even having different topologies (T level) and secondary structure architecture (A level). However, their active/binding sites and overall conformation of the catalytic core occasionally retain structural similarity.

This is exemplified in two mammalian tyrosine phosphatases (EC 3.1.3.48) that have a remarkably similar catalytic core but completely different overall folds (CATH 3.40.50.2300 for the bovine low molecular weight phosphatase, vs. CATH 3.90.190.10 for the human tyrosine phosphatase). For these two analogues, Gherardini et al. suggest convergence[3], however, the striking similarities in the vicinity of the active site could infer two additional plausible scenarios: a) divergence to accommodate different metabolic needs and subcellular location (endoplasmic reticulum vs. cytoplasm), with conservation of the catalytic core, or b) divergence towards a different function, followed by re-convergence to the initial one. Another example of evolutionary ambiguity is the pair of two bacterial/mammalian 3-hydroxydecanoyl dehydratases (EC 4.2.1.59) that share high similarity in the extended active site, with four catalytic residues superposing at very low RMSD (0.62Å). This is accompanied by high mechanistic similarity. Our observations lead us to believe these analogues are either an extreme case of convergent evolution, or a case of horizontal gene transfer from symbiotic bacteria to the host, followed by widespread structural divergence, retaining the structure of the first residue shells.

Evolutionary ambiguities are expected to be common in activities that are: a) metabolically ubiquitous –i.e. found in multiple different organisms and metabolic pathways, and b) mechanistically specific –i.e. demanding a specific chemical toolkit and catalytic steps to happen.

### Limitations of EC classification

We identified enzymes of similar EC classification that have substantial functional differences. Although we are looking at identity in all 4 EC levels, which should imply identity in substrate selectivity, this is fuzzy in some enzymes, such us polymer-acting ones, or ones using multiple substrates, like chloride peroxidases (EC 1.11.1.10). Three isozymes on our set share this EC and are totally unrelated. Structures, mechanisms, and metabolic contexts are completely different, and atheir only commonality is selectivity for Cl^-^ ions and H_2_O_2_. The third cognate substrate is a variable organic compound that is chlorinated.

Reactions like this are generic and involve several components, one of which is simplistic (like an ion). Also, Markush structures are often used to represent ensembles of substrates that share a common ‘catalysable’ moiety. In such cases, EC often classifies the enzyme based on the simplest cognate ligand. This means that functional differences could be concealed by the EC classification process. Similar classification ambiguities are observed in promiscuous enzymes, such as proteases or DNAses, leading to the same problem. Additional complexity in EC classification can derive from the type of cognate ligand considered for defining the 4^th^ number.

In most hydrolases, for example phosphatases from the 3.1.3.x sub-subclass, the sub-sub-subclass is defined by substrate selectivity whereas in triterpene synthases (5.4.99.x), it is defined by product selectivity.

## Discussion

We have collected a sample of isozymes with sufficient structural and functional evidence, to reveal some “rules” or paradigms of convergent evolution. These data, albeit limited, have allowed us to identify cases of chemical convergence in enzymes (catalysis of the same overall reaction), accompanied by structural convergence in the active site and mechanistic convergence. We showed that these three levels of similarities might not necessarily co-exist, and based on this, we defined three paradigms of convergence: structural and mechanistic (13 cases), mechanistic-only (11 cases) and reaction-only (9 cases).

Our methodology for comparing isozymes, among others, included examination of cofactor selectivity. If we investigate the paradigms further by considering cofactor selectivity, there were 4 unobserved sub-paradigms in this survey. One sub-paradigm refers to enzymes with similar active site geometries, dissimilar mechanisms, and the same cofactor selectivity. This might also not be observed because having these traits in common is usually indicative of a homologous relationship. Conversely, we did not observe any enzymes that had similar active site geometry and mechanism, but significantly different cofactors (e.g. a metal vs. organic cofactor); this is also expected as the presence of cofactors would affect the geometry of the active site. We also do not see a pair of enzymes with similar active site geometries and mechanisms, where one enzyme uses a cofactor, and one does not. This is also expected for the same reason as the previous sub-paradigm; however, we observe this when mechanisms differ (e.g. in acid phosphatases). Although there are examples where a reaction can be performed under both a cofactor dependent and independent mechanism, these are mostly classified into the “reaction-only” paradigm. This is probably because it is quite unlikely that enzymes would have the same geometry and cofactors, without having any mechanistic commonality. Overall, these results could imply some level of rigidity that comes with active site geometries being similar. Last, we do not see any cases where two enzymes have significantly different cofactors that are both present, and similar mechanisms. This could be attributed to the limited variety of cofactors in nature, where each cofactor has evolved to serve a specific role in catalysis[84]; thus, it is unlikely for two enzymes to evolve to bind different cofactors and still have a similar mechanism.

Classification of isoenzyme groups into paradigms was performed in a semi-manual way, by calculating some quantitative measures of similarity (catalytic residue superposition, mechanistic similarity), and by referring to available literature. In this process, we realised that the quantitative measures alone are not sufficient for assignment into paradigms, therefore, each case was examined individually and, if necessary, the quantitative results were overridden. For example, there were several cases where geometrical similarities were found to be spurious, and the active sites had to be examined manually to ensure biological relevance. Interestingly in the context of template matching: we have found that smaller templates, consisting of a few atoms, may yield spurious matches, even if the search is restricted to active sites. Therefore, one needs to be aware of false positive results in template-based searches, especially when these are performed at a high-throughput level.

To address the question of why solutions to chemical functions have evolved several times, we have discussed several examples of comparable chemical reactions by analogous enzymes. These data indicate that the factors influencing convergent chemistry between non-homologues enzymes are: 1) a common need for a metabolic reaction, but independent evolution has led to the emergence of different molecules. This is mostly observable in analogues from different kingdoms of life. 2) a different metabolic context demanding different enzymes. 3) different regulation of isoenzymes leading to enzymes of functional promiscuity. 4) Environmental pressures to solve biological challenges with the limited raw material available (i.e. amino acids, water, metals, etc.) lead to similar solutions, with the more efficient being retained.

Analysing how proteins with completely different folds maintain identical catalytic activity, we found many cases where a reaction might be mechanistically and/or structurally constrained. The more constrained the process, the higher the similarities between analogues (e.g. in phosphatases and in proteases). Other processes are less constrained, and convergence is only observable at the end of the reaction. Our approach to examining convergence from various points of view was hybrid, including both manual inspection and automated/quantitative methods. The latter can be scaled in a reproducible pipeline, allowing a more systematic and comprehensive analysis of convergence in the future, not necessarily restricted within the limits of M-CSA. In the interest of clarity and conceptualisation we have generalised the observations from the data, where each of the pairs of enzymes evaluated provides unique insights into how proteins can maintain the same catalytic potential on a completely different overall shape.

## Methods

### Data Preparation

Groups of enzymes catalysing the same reaction and having different folds were collected from M-CSA[85] using the public API, by querying for entries with identical EC numbers and different CATH[86] numbers in the catalytic domain of their reference structure. Entries were grouped by EC number, and those groups having redundant CATH numbers, multiple CATH numbers (e.g. active site formed in the interface of two domains), or incomplete information (not all 4 EC levels determined, no CATH mapping, etc) were removed. In cases of EC groups having more than two enzymes after filtering, relationships were analysed in pairs, so that groups with 3 enzymes would result in 3 distinct analyses. This first filtering resulted in 46 pairs of enzymes that perform the same function and have different folds, across 34 unique EC numbers. Some extra filtering was done to ensure that this survey focuses solely on pairs that have similarities due to functional convergence. From the original set of 46 pairs, 4 were excluded as sequence or structural similarity implied potential homology. Pairs from the restriction enzyme group (EC 3.1.21.4, accounting for six pairs) were also excluded foe the same reason [67–69]. 5 pairs were taken out due to limited available information. In several cases, two enzymes may not bind identical substrates or products, but those are still in scope since some of them are substrate promiscuous. Most of these were due to unspecified -R groups, different substrate specificities, or both enzymes manipulating the same functional groups on different molecules. The final, reduced set included 31 pairs across 23 EC numbers. 21 of the 31 pairs were found to have mechanism similarity, however none of the pairs showed complete identity.

### Literature analysis

Each pair was analysed using information found on M-CSA (enzyme mechanism and catalytic residue reaction roles), PDBe[87] (representative crystal structure selection, assembly composition), PDBsum[88] (bound ligand information), and the enzymes associated literature from these databases.

### Sequence and structure comparison

Local and global sequence alignments were performed using the Needleman-Wunsch and Smith-Waterman algorithms respectively via the EMBOSS service[88,89]. Alignments were visualized in JalView[90].

### Active site structure comparison

Functional atom templates[9] (three atoms per residue) were generated for each active site, taking all possible combinations of 2, 3 and 4 residues, using the CSA-3D package of our previous work[91]. Template definitions are mechanism-informed (e.g. three backbone atoms are selected for residues known to contribute their backbone to the mechanism) and allow for alternative matching of chemically similar atoms and residues, as described in ref. [6]. Templates of the same size were structurally compared using the template matching program Jess[92], in an “all against all” fashion. The match with the lowest RMSD and highest number of residues was selected as a reference atom-atom mapping frame for dynamic active site fitting.

### Dynamic active site fitting

Using the active site atom-atom mapping method described above, enzyme pairs were fitted on their functional atoms, using a Gaussian-weighted version of the Kabsch algorithm[93,94], as described in ref. [91]. The algorithm outputs the two enzymes with their coordinates transformed accordingly, along with a weighted RMSD (wRMSD), an unweighted RMSD and a coverage value. The latter corresponds to the proportion of mapped catalytic residues over the number of residues of the largest of the two active sites.

### Catalytic mechanism comparison

Mechanism comparison was conducted, which resulted in pairs being categorised as having “similar” or “not similar” mechanisms which was determined by seeing if the enzymes had *any* similarities in their mechanism. Similarities included near identical steps (ex: nucleophilic attack on the same atom, not necessarily the same nucleophile) or identical order of some or all events. This was initially done by manual inspection and comparison of the mechanisms found on M-CSA, and in cases where both enzymes in a pair had detailed mechanism description, a mechanism similarity score was calculated as follows: the data on the curly arrow diagrams of each catalytic step are first abstracted into a set of curly arrows and their chemical environments. This representation of each curly arrow is called an arrow-environment and it includes the atoms interacting directly with the curly arrow and two shells of atoms and bonds around those. The similarity score for each mechanism pair is then calculated as the Jaccard index of the two sets of arrow-environments (*Ribeiro et al.* [95] *in preparation*).

### Cofactor binding comparison

Cofactor preferences were compared for each enzyme pair, focusing only on cofactors directly involved in catalysis. The result of this analysis was placement into two broad categories that addressed identity, with pairs having the same cofactors being marked “yes” and ones with different cofactors being marked “no”. Within the “yes” and “no” categories there were subcategories. In the “yes” category pairs were marked as both having present cofactors that were identical, or both having no cofactor present. In the “no” category pairs were marked as both having cofactors that are different, or one having a cofactor present and one not having a cofactor present. We considered NAD and CoA cofactors if they were in the reaction regardless if their original chemical composition was regenerated at the end of the catalytic cycle.

## Acknowledgements

The work was supported by the EMBL International PhD Programme (IGR) and the European Molecular Biology Laboratory (JCK, GO, AJMR, NB, JMT)

## Author contributions

**IGR**: Conceptualisation, Methodology, Software, Validation, Writing – Original Draft preparation

**JCK**: Methodology, Investigation, Visualisation

**GO**: Software, Methodology

**AJMR**: Data Curation, Validation

**NB**: Conceptualisation, Validation, Writing – Review & Editing

**JMT**: Conceptualisation, Supervision, Resources, Funding Acquisition, Project Administration, Writing – Review & Editing

